# Precise measurement of molecular phenotypes with barcode-based CRISPRi systems

**DOI:** 10.1101/2024.06.21.600132

**Authors:** Joseph H. Lobel, Nicholas T. Ingolia

## Abstract

Genome-wide CRISPR-Cas9 screens have untangled regulatory networks and revealed the genetic underpinnings of diverse biological processes. Their success relies on experimental designs that interrogate specific molecular phenotypes and distinguish key regulators from background effects. Here, we realize these goals with a generalizable platform for CRISPR interference with barcoded expression reporter sequencing (CiBER-seq) that dramatically improves the sensitivity and scope of genome-wide screens. We systematically address technical factors that distort phenotypic measurements by normalizing expression reporters against closely-matched control promoters, integrated together into the genome at single copy. To test our ability to capture post-transcriptional and post-translational regulation through sequencing, we screened for genes that affected nonsense-mediated mRNA decay and Doa10-mediated cytosolic protein decay. Our optimized CiBER-seq screens accurately capture the known components of well-studied RNA and protein quality control pathways with minimal background. These results demonstrate the precision and versatility of CiBER-seq for dissecting the genetic networks controlling cellular behaviors.

## Introduction

Comprehensive, genome-scale reverse genetic screens have successfully uncovered the molecular underpinnings of diverse biological processes^1^. CRISPR-Cas9 is a versatile tool for reverse genetics that can be targeted by guide RNAs (gRNAs) to create programable genome-wide knockdowns or knockouts^2–4^. By determining which guides alter a specific cellular behavior, one can identify the responsible regulatory genes and prioritize these for mechanistic studies. These approaches have been widely used, and increasing the sensitivity and range of phenotypes accessible to these platforms would have broad benefits^1^.

A central challenge in CRISPR-based reverse genetics is coupling genetic perturbations to their biological effects on a genome-wide scale. Towards this end, our lab combined <u>C</u>RISPR interference (CRISPRi) with <u>B</u>arcoded <u>E</u>xpression <u>R</u>eporter <u>seq</u>uencing (CiBER-seq) to directly and quantitatively measure diverse molecular phenotypes by high-throughput sequencing^5^. CiBER-seq uses a library of gRNAs that are linked with a transcribed reporter containing a unique, guide-specific barcode; guides that change expression of the reporter—for example, by perturbing a regulator—will change expression of their individual, guide-specific barcode. Measuring reporter expression changes by deep sequencing of the barcodes allows for simple, pooled, genome-wide interrogation of numerous biological processes^5,6^. Reporter expression can assess many nuanced phenotypes that are inaccessible by simply measuring cell growth and survival. By quantifying reporter expression through deep sequencing, CiBER-seq also offers substantial advantages compared to fluorescent protein reporters analyzed by flow sorting and sequencing, which discretizes effects into specific bins and has limited throughput. More generally, RNA reporters are better equipped to directly address the regulation of RNA metabolism and post-transcriptional processes^5,7^. Thus, coupling CRISPRi with barcode measurements enables quantitative reverse genetic screens across a broader range of phenotypes and cellular conditions.

While CiBER-seq and other barcode-based CRISPRi systems have emerged as powerful tools to assess genotype-to-phenotype relationships, these approaches are still in their infancy^5,7^. In any genetic screen, high background signal can obscure interesting candidates and prevent identification of critical regulatory factors. This background can arise when perturbations affect technical aspects of the reporter system rather than the biological process under investigation, leading to the spurious appearance of phenotypic effects. Technical factors can also add noise or distort measurements of these effects. For example, sequencing-based genetic screens use combinations of RNA and DNA barcodes with different handling and processing steps that can lead to variation in barcode measurements^8^. Overcoming these technical and systems-level effects would enhance the ability to quantitatively define regulators of cellular responses.

Here, we dramatically improve the sensitivity of barcode expression reporters and expand their scope to interrogate diverse molecular processes. We essentially eliminate background in CiBER-seq experiments by expressing RNA barcodes from two closely matched promoters and further reduce noise through precise, single-copy integration of these reporters. This generalizable approach enables accurate dissection of genetic networks controlling diverse protein and RNA-level phenotypes. We use this system first to profile genetic regulators of a potent protein degradation signal, finding the established ubiquitin-proteasome system known to drive its turnover. We then adapt CiBER-seq to examine mRNA quality control directly and identify the conserved nonsense-mediate decay factors that are required to degrade a transcript containing a premature termination codon. In both cases, our optimized platform finds known regulators with exquisite precision and minimal background. These simple and effective improvements for barcode-based CRISPRi screens are readily adaptable to investigate genetic regulators across a range of molecular phenotypes.

## Results

### RNA barcodes expressed from matched promoters eliminates background in barcode-based CRISPRi screens

CiBER-seq is a massively parallel reporter approach that measures how CRISPRi-based gene knockdown affects a transcriptional reporter through deep sequencing of an expressed mRNA barcode^5^. These barcodes, embedded within the 3′ UTR of a reporter transcript, are uniquely linked to a guide on the same plasmid **(Fig. 1A)**^9^. Comparing the abundance of each expressed RNA barcode before and after induction of guides reveals how the CRISPRi knockdown of a gene influences expression of its associated reporter **(Fig. 1B)**. Accurate expression measurements require normalization of barcode RNA abundance to correct for the abundance of cells containing that barcode within the population, which varies between experiments and can change due to growth defects. Typically, DNA barcodes—which should be present at one copy per cell—are sequenced to control for these variations **(Fig. 1B)**. RNA-to-DNA ratios have been used in previous CiBER-seq screens by our lab and in other barcode-based CRISPRi screens to interrogate diverse biological regulatory programs^5,7^. While these methodologies have yielded valuable discoveries, improving the precision and scope of these technologies promises to more accurately decipher genetic networks underlying complex cellular processes.

**Figure 1:**
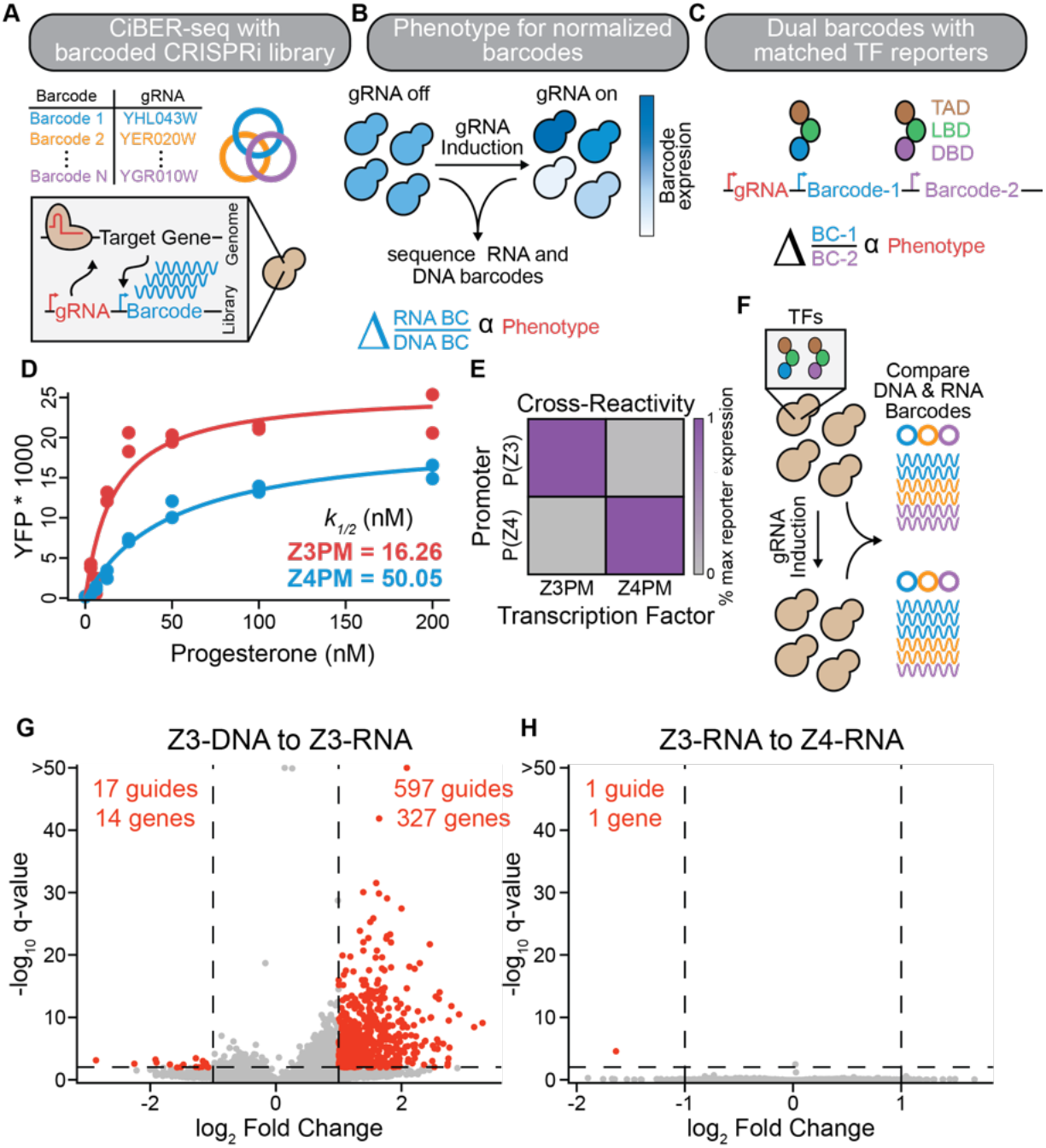
Eliminating background in barcode-based genetic screens. **(A)** Schematic of paired guide-barcode libraries for CRISPRi screening. **(B)** Workflow for CiBER-seq screen and determination of phenotypic effects. Sequencing libraries are prepared from samples before and after guide induction, normalizing RNA barcodes to DNA barcodes. **(C)** An idealized CiBER-seq platform using closely matched transcription factors that each express a barcode from similar promoters with common genetic dependencies. **(D)** Characterization of the dose response of each hormone-inducible transcription factor by flow cytometry of yeast transformed with a YFP expressed from the cognate promoter. Mean YFP was fit to a simple binding isotherm for biological replicates, see Methods (n=2). **(E)** Cross reactivity of Z3PM or Z4PM expressing YFP from a P(Z3) or P(Z4) promoter. YFP expression is normalized to the transcription factor and its cognate promoter (n=2). **(F)** Schematic for evaluating technical variations between DNA and RNA barcode-based comparisons. All barcodes are isolated from the same sample before and after guide induction. **(G)** Analysis of genome-wide CiBER-seq screen with Z3PM RNA barcodes normalized to DNA barcodes levels. Each point is a single guide, with significant guides colored red. A q-value < 0.01 and > 1 log_2_-fold change and is represented by dashed lines. **(H)** Same screen as in **(G)**, except Z3PM barcode expression was normalized to the control barcodes driven by Z4PM.

A key challenge in any high-throughput screen is distinguishing the biological phenotype of interest from effects caused by technical aspects of the system. In massively parallel reporter assays, differences in sample preparation between RNA and DNA libraries can distort expression estimates. Additionally, in CiBER-seq, guides that affect core cellular processes may influence barcode DNA abundance or RNA expression, leading to biologically irrelevant changes in the measured RNA-to-DNA ratio. Collectively, these technical differences can cause high background that masks sought-after targets and necessitates laborious curation and validation of individual candidates. To mitigate these effects, previous work from our lab expressed reporter and normalizer barcodes from two different promoters. These RNA-to-RNA comparisons controlled for many non-specific guide effects and eliminated technical differences that arise from separately handling plasmid DNA and mRNA. Nonetheless, we found that different genetic requirements of promoters substantially altered the ratio of reporter to normalizer barcodes; hundreds of guides had activity by affecting general transcription machinery rather than the phenotype of interest^5^. Correcting for these systems-level effects would increase the sensitivity and robustness of CiBER-seq.

We reasoned that barcodes expressed from closely matched promoters would share the same genetic dependencies and measuring the ratio between these barcodes should eliminate much of the systematic and technical variation in CiBER-seq **(Fig. 1C)**. Achieving this idealized setup requires promoters that respond to biologically interesting regulation while otherwise sharing similar genetic requirements. We turned to synthetic transcriptional reporters driven by chimeric transcription factors that exclusively bind their cognate synthetic promoter. We used two different promoters derived from the same core *GAL1* sequence that differ only in the binding sites for distinct, heterologous zinc-finger DNA binding domains, termed “Z3” and “Z4”. These DNA-binding domains are incorporated into artificial hormone inducible transcription factors, Z3PM and Z4PM, that share a common progesterone binding and Msn2 *trans*-activation domains^10,11^ **(Fig**.**1 C)**. We also appended a PEST degron domain to increase transcription factor turnover and ensure protein levels more accurately track the short half-lives of mRNAs in yeast^12,13^. We confirmed that these transcription factors had similar hormone responses, are highly specific for their cognate promoter, and show no growth defects when induced, making them ideal candidates to drive barcode expression in CiBER-seq **(Fig. 1D/E & Fig. S1A-D)**.

We assessed how comparing barcodes expressed from these closely matched reporters could minimize background effects seen in previous iterations of CiBER-seq. As a test case, we performed a screen using reporter and normalizer barcodes driven by the Z3PM or Z4PM transcription factors, respectively, with no additional differences between them, and carried out deep sequencing of both DNA and RNA barcodes from the same samples **(Fig. 1F)**. Given the strong similarity between these reporters, few guides should cause significant changes and any active guides could be attributed to systematic background. We transformed yeast expressing both Z3PM and Z4PM with a plasmid-based guide library containing barcoded reporters driven by these transcription factors, grew cells at a constant density, and collected cells before and after ∼9 h of guide induction (∼6 doublings)^14^. We isolated RNA and DNA in parallel, counted barcodes by deep sequencing, and identified guides that caused significant barcode changes using established linear models for massively parallel reporter assays^15^. Knockdown of hundreds of genes changed the RNA-to-DNA barcode ratio; many were involved in DNA replication or RNA metabolism **(Fig. 1G & S2)**. In stark contrast, comparing Z3- and Z4-driven RNA barcodes revealed only one significantly different guide **(Fig. 1H)**. Since these comparisons were made using the exact same biological samples, we conclude that technical effects distort RNA-to-DNA measurements, increasing background that would obscure guides driving the phenotype of interest. Both comparisons had >14 million reads per sample, and read counts were within ∼2-fold, ensuring these differences represent biological effects and are not due to sequencing depth or statistical power. Closely matched orthogonal promoters, driven by synthetic transcription factors, provide a simple solution to dramatically reduce background in CiBER-seq and similar approaches, and should thereby enable more precise identification of relevant targets in reverse genetic screens.

### High-efficiency genomic integration of CiBER-seq libraries

Like many other massively parallel experiments conducted in yeast, CiBER-seq relies on libraries expressed from autonomously replicating plasmids. Plasmid loss and copy number variation, arising due to imperfect plasmid segregation, introduces technical noise into reporter measurements^16,17^. Furthermore, continuous selection is required to maintain plasmids and constrains the growth conditions that can be used. While genomic integration solves these limitations, homology-directed integration into the yeast genome occurs at much lower efficiency than plasmid transformation; this bottleneck limits its application in massively parallel experiments where large numbers of transformants are needed to span diverse libraries. Double-stranded breaks created by CRISPR-Cas9 or other nucleases can enhance integration, but this process is not benign and remains too inefficient for use with large-scale libraries^17^. In human cells, serine recombinases, such as Bxb1, have been utilized to create genomic landing pads that support high-efficiency integration of complex libraries^18–20^. Bxb1 works efficiently in yeast, and we reasoned that a similar landing pad might allow high-diversity, chromosomally integrated libraries in CiBER-seq screens^21^.

We created CiBER-seq libraries of donor plasmids with guide RNAs and barcoded reporters that could be integrated into a single genomic landing pad **(Fig. 2A)**. We drew from other systems to provide streamlined integration and stringent selection of transformants. Specifically, our landing pad constitutively expresses a yeast-optimized Bxb1 recombinase gene that is inactivated by integration of a donor plasmid, eliminating the need for additional components or recombinase induction. Integration of the donor plasmid reconstitutes a *URA3* selection marker gene from two halves that are split between the landing pad and the donor, ensuring marker expression only upon integration specifically at the landing pad^22,23^. We see no background growth in selective conditions in the absence of on-target integration, consistent with previous studies using similar split markers.

**Figure 2:**
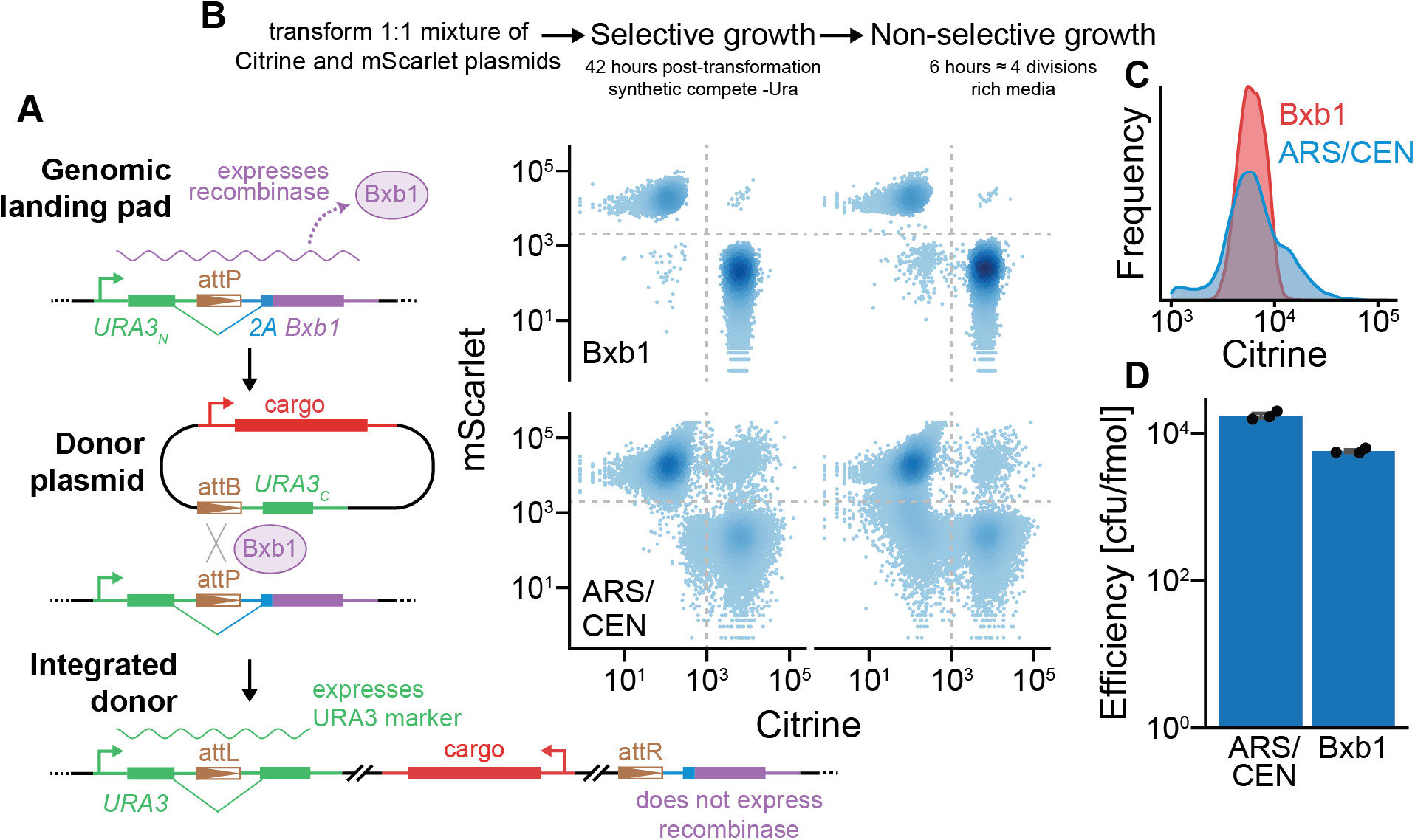
High-efficiency, single-copy integration of reporters with a Bxb1 recombinase-based system. **(A)** Schematic of yeast-based Bxb1 integration system. Bxb1 is constitutively expressed until recombined with a donor plasmid, which reconstitutes a *URA3* selectable marker. **(B)** Flow cytometry of yeast transformed with a mixture of plasmids encoding yECitrine or yEmScarlet through plasmid-based or Bxb1-mediated recombination approaches. Fluorescence was recorded both before and after removal of selective pressure. **(C)** Distribution of yECitrine fluorescence of yeast transform through different approaches **(D)** Transformation efficiency of plasmid-based or Bxb1-mediated recombination approaches (n=3).

We then tested whether landing pad integration resolved the sources of noise seen in autonomous plasmids. We transformed yeast with an equal mixture of plasmids expressing the yellow fluorescent protein yECitrine and the red fluorescent protein yEmScarlet-I to compare autonomous plasmids with Bxb1 donors **(Fig. 2B)**. We carried out bulk selection in liquid culture and then analyzed the transformed populations by flow cytometry. After selection, virtually all yeast transformed with the Bxb1 donor exclusively expressed one fluorescent protein, which persisted for several generations after selection was removed. In contrast, plasmid transformation yielded a large population of cells that lacked plasmid and quickly expanded when selection was relaxed. Furthermore, a substantial number of yeast expressed both fluorescent proteins simultaneously, indicating the presence of multiple, distinct plasmids within the cell. In the context of a complex library transformation, such as a CiBER-seq experiment, yeast containing two different library plasmids would contribute to technical noise. This difference in plasmid levels also produced more variation in fluorescence protein expression than Bxb1-mediated integration **(Fig. 2C)**. Importantly, Bxb1-mediated integration was efficient enough for large-scale transformation of complex libraries and was within ∼3-fold of autonomous plasmids transformation **(Fig. 2D)**. Thus, recombinase-mediated integration at a genomic landing pad ameliorates several technical challenges posed by plasmid library transformations while maintaining high efficiency required for large-scale genetic screens.

### Optimized CiBER-seq identifies regulators of a degron

We aimed to demonstrate how this optimized CiBER-seq platform could characterize diverse molecular phenotypes with high precision. We first surveyed genetic modifiers of targeted protein degradation by assessing regulators of this pathway through a matched-promoter CiBER-seq library integrated by Bxb1-mediated recombination. We appended the well-studied CL1 degron to the Z3PM transcription factor to destabilize it and thereby decrease Z3-driven barcode expression **(Fig. 3A)**. Indeed, Z3 barcode expression was ∼40x lower than Z4 barcodes driven by the untagged Z4PM construct. The CL1 degron is ubiquitinated by the E3 ligase Doa10 and subsequently delivered by Cdc48 to the 26S proteasome for degradation^24,25^. Reduced Z3PM barcode expression was fully rescued when Doa10 was knocked out, confirming that our reporter detects genetic changes in CL1 degradation **(Fig. 3B)**. This strong protein-level phenotype was well suited to interrogate with our optimized CiBER-seq approach.

**Figure 3:**
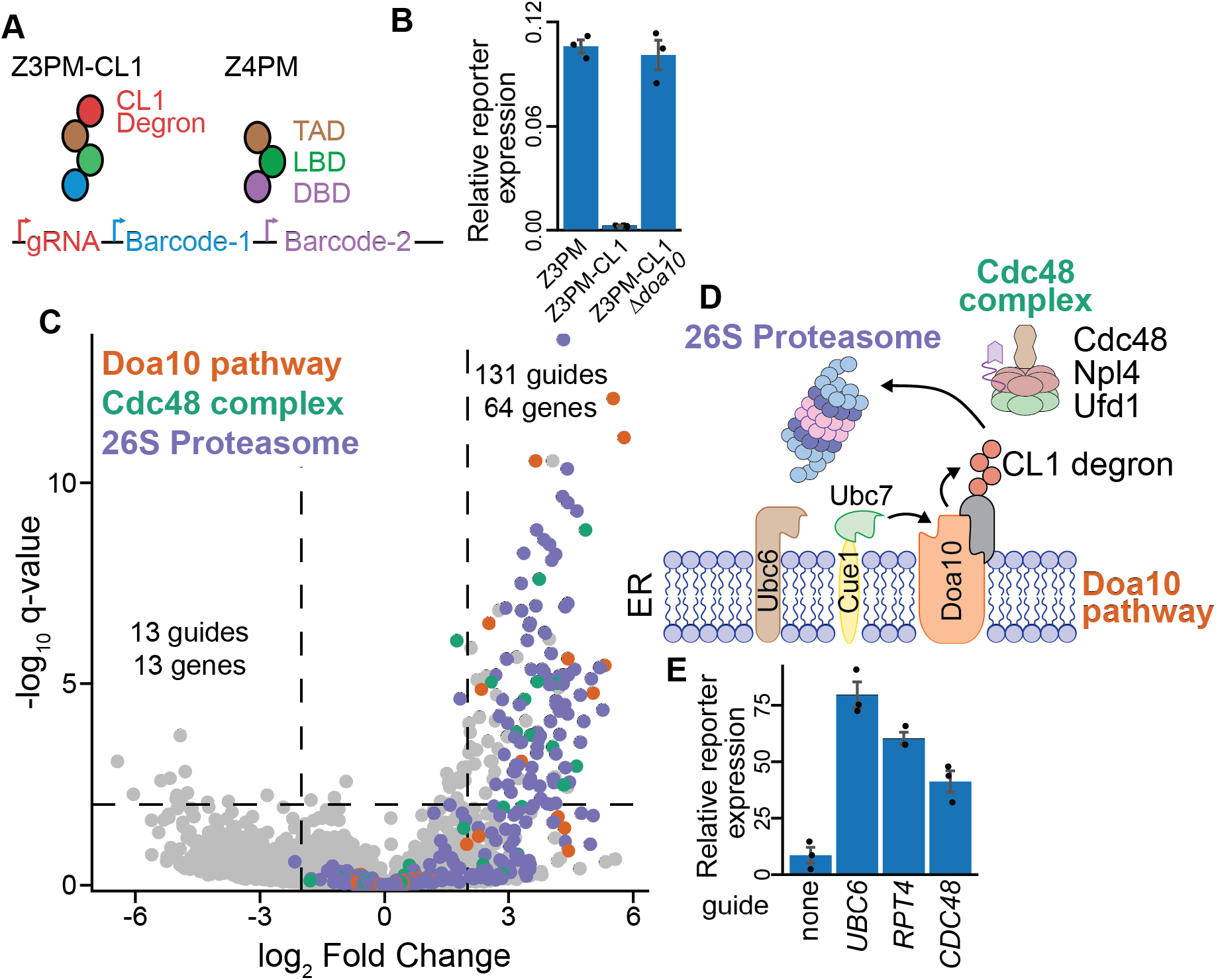
Precise identification of degron regulators with optimized CiBER-seq. **(A)** Schematic for CiBER-seq to characterize regulators of the CL1 degron. **(B)** RT-qPCR of reporter transcripts expressed by Z3PM or Z3PM-CL1 in a wildtype or *Δdoa10* background, compared to the normalizer transcript (n=3). **(C)** Analysis of genome-wide CiBER-seq screen for regulators of the CL1 degron. Each point is a single guide and colored based on molecular function in legend. Significant and robust guides were assessed by a q-value < 0.01 and > 2 log_2_-fold change, which is represented by dashed lines. **(D)** Schematic of established CL1 turnover pathway with yeast protein names displayed. **(E)** RT-qPCR of reporter barcodes expressed by Z3PM-CL1 with guides induced (n=3).

We then performed a genome-wide CiBER-seq screen to characterize regulators controlling the stability of the CL1 degron. We integrated a CiBER-seq dual-barcoded guide library using Bxb1-mediated recombination into a reporter yeast expressing both Z3PM-CL1 and Z4PM and quantified changes in barcode ratios after guide induction. As expected, many guides that stabilized the reporter transcription factor corresponded to genes found to be involved in turnover of the CL1 degron through traditional knockout or temperature-sensitive alleles **(Fig. 3C/D)**. For example, we identified all components of the Doa10 E3 ligase system, including the sequential E2 enzymes Ubc6 and Ubc7 along with the Ubc7 activator, Cue1^26^. We also identified many guides targeting the components of the 26S proteasome and members of the Cdc48 complex **(Fig. 3D)**. Overall, this approach identified 131 guides targeting 64 genes that stabilized the reporter transcription factor, ∼78% of which were involved in either the 26S proteasome, Doa10 ubiquitination, or Cdc48 pathways, demonstrating the high precision of our optimized CiBER-seq approach^24^. In contrast, guides that appeared to further destabilize the Z3PM-CL1 degron likely represent a low background, as none of these targeted the same gene, were minimally above our statistical cutoff, and had no enriched gene ontology terms **(Fig. 3D)**.

A major advantage of CRISPRi is its ability to target essential genes without relying on temperature-sensitive alleles. For example, we found numerous guides targeting different components of the essential 26S proteasome or Cdc48 complex. We individually validated several of these guides targeting both essential genes and non-essential and components of the CL1 degradation pathway. Knockdown of these genes increased barcode levels expressed by the Z3PM-CL1 construct, consistent with stabilization of the reporter transcription factor **(Fig. 3E)**. These results demonstrate how an optimized CiBER-seq platform can easily interrogate regulators of specific phenotypes, such as protein turnover, with exquisite specificity and sensitivity.

### Expanding optimized CiBER-seq to characterize regulators of RNA phenotypes

Given the success of our optimized CiBER-seq to confidently identify the full pathway responsible for CL1 degradation, we next wanted to further expand the phenotypes we could address with this system. We reasoned we could extend CiBER-seq to examine post-transcriptional regulatory processes, which shape gene expression by modulating mRNA half-lives or degrading faulty messages. Despite their central role in cellular processes, RNA-level phenotypes are currently only indirectly measured through standard genetic screens. Typical studies couple problematic mRNAs to expression of a fluorescent protein, which is used as a proxy for transcript abundance^27–33^. While valuable, these systems do not distinguish the separate contributions of translational control and transcript stability to the observed response. Our understanding of these complex post-transcriptional regulatory programs would be aided by methods that can directly interrogate regulators of RNA-level processes.

CiBER-seq is well positioned to break this barrier because it infers guide effects by measuring RNA barcodes. Inserting barcodes within the 3′ UTR of a problematic mRNA would allow a genome-wide CRISPRi screen that directly assesses how each guide influences levels of the reporter transcript **(Fig. 4A)**. As a test case, we chose to characterize regulators of nonsense mediated decay (NMD), a translation-coupled process that triggers destruction of messages harboring a premature termination codon (PTC). During NMD, release factors recognize the PTC within the ribosome and assemble Upf proteins, which recruit the decapping machinery to trigger degradation of the message **(Fig. 4B)**^34^. To profile regulatory factors that control turnover of a NMD substrate, we designed a reporter mRNA that possessed a PTC and a normalizer that lacked one. We constructed a CiBER-seq library that embedded the barcodes within the 3′ UTRs of the NMD and control reporters, where they serve as a direct measurement of mRNA levels. As before, we sought to minimize background by ensuring both transcripts were expressed from matched promoters **(Fig. 4A)**. We then performed a genome-wide CiBER-seq screen by sequencing barcodes from samples taken before and after guide induction. Unlike protein-based reporters, direct measurements of RNA allowed us to additionally interrogate phenotypic changes that occur when translation is arrested. Because a hallmark of NMD is its dependence on active translation, we assessed this requirement by also collecting barcodes 1 hour after treating cells with cycloheximide, which stabilized the PTC containing reporter **(Fig. 4C/D)**^35^.

**Figure 4:**
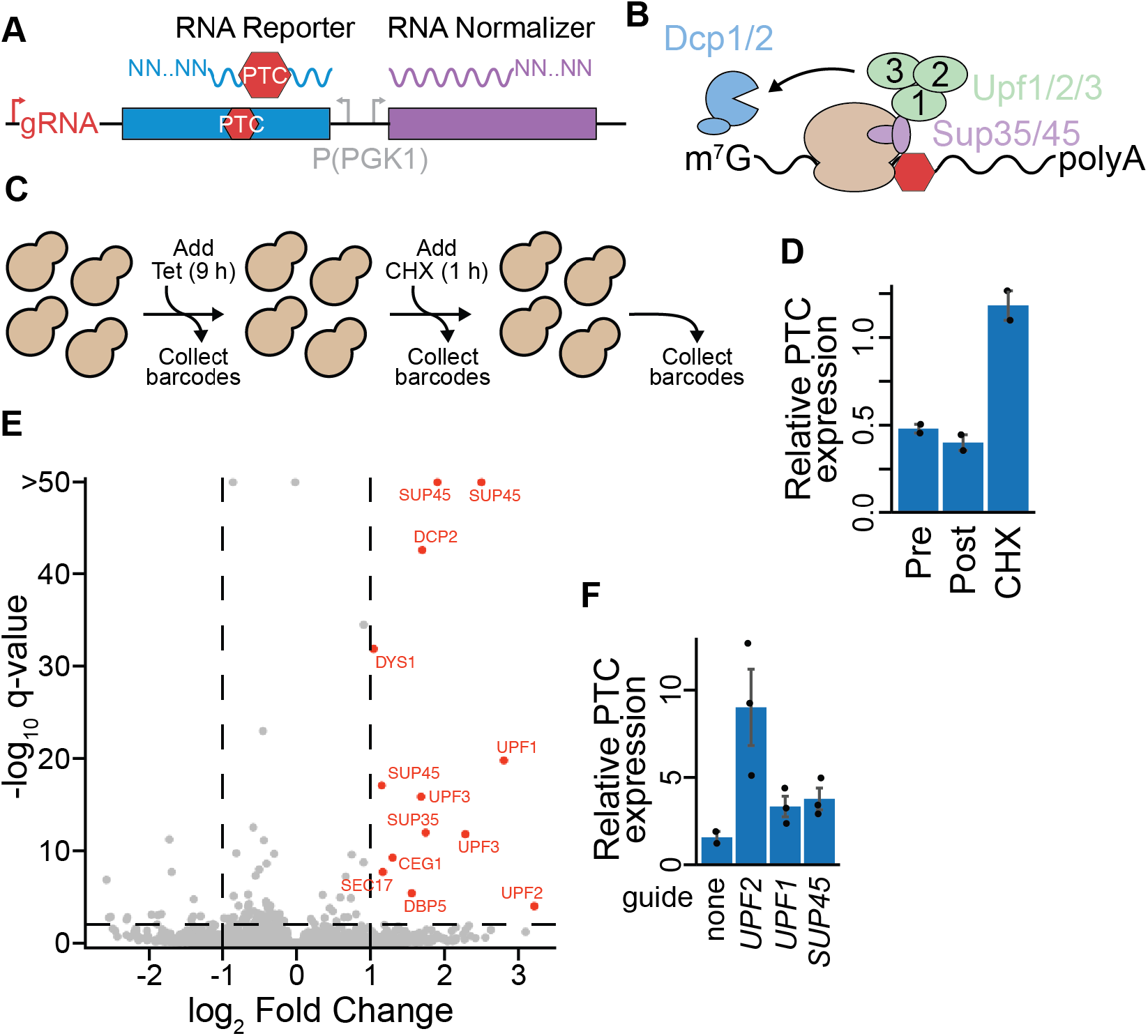
CiBER-seq directly measures regulators of an RNA quality control pathway. **(A)** Schematic for using CiBER-seq to interrogate RNA-level phenotypes. Barcodes are embedded in the 3′ UTR of a reporter and normalizer, which are constitutively expressed from identical promoters. **(B)** Proteins involved in NMD, with yeast gene names displayed. **(C)** Workflow for CiBER-seq to investigate regulators of NMD and their dependency on active translation. **(D)** RT-qPCR of the NMD reporter, compared to the normalizer transcript without the PTC, at all steps during CiBER-seq (n=2). **(E)** Analysis of genome-wide CiBER-seq screen for regulators of NMD. Guides with a q-value < 0.01 and > +1 log_2_-fold change are labelled in red, with thresholds represented by dashed lines. **(F)** RT-qPCR of PTC-containing mRNA compared to normalizer with guides induced (n=2-3).

Again, our optimized CiBER-seq screen precisely identified key NMD genes with minimal background. We found strong NMD suppression from guides targeting both Sup35 and Sup45 (which encode the release factors eRF1 and eRF3), the three Upf proteins, and the decapping protein Dcp2 **(Fig. 4B/E)**. More broadly, these guides were strongly enriched for NMD and translation termination gene ontology terms **(Fig. S3A)**. Indeed, 10 out of the 13 guides that stabilized reporter expression targeted genes with known roles in NMD, while guides that appeared to further destabilized the PTC reporter did not share any common genes and lacked any significant gene ontology terms. Experiments performed with individual guides confirmed that these knockdowns stabilized the PTC reporter and recapitulated the rank order of phenotypes observed in our CiBER-seq screen **(Fig. 4F)**. These factors lost activity when translation was arrested with cycloheximide, consistent with the requirement of ribosome elongation for NMD **(Fig. S3B/C)**^34^. This work highlights how CiBER-seq can uncover molecular phenotypes in the absence of ongoing translation, something not possible with fluorescent protein-based or survival screens. We again observed minimal background effects in this distinct screen, revealing the general applicability of using matched promoters for eliminating background in genome-wide barcode-based CRISPRi platforms **(Fig. 4E)**. Combining our optimized CiBER-seq with direct measurements of RNA reporters enables CRISPRi approaches to quantitatively and precisely interrogate diverse RNA as well as protein level molecular phenotypes.

## Discussion

The combination of CRISPRi genetic perturbations and barcoded expression reporters holds clear promise as a widely applicable tool for biological discovery. Increasing the range of processes that can be assessed with barcoded reporters and reducing noise in these measurements would further enhance these powerful techniques. Here, we realize the full potential of CiBER-seq platforms by addressing the major sources of background with simple and generalizable modifications. We find that using RNA reporter and normalizer barcodes, driven from closely matched promoters, eliminates irrelevant technical effects across multiple screens. Recombinase-mediated integration further ensures stringent isolation of single guides in each cell. We combine these elements in CiBER-seq screens that accurately and comprehensively identify known regulators in post-transcriptional and post-translational control pathways. These results illustrate how this optimized CiBER-seq strategy provides a broadly-applicable platform for genome-wide screens.

While genome-wide knockout or knockdown approaches have been invaluable for mapping regulatory networks, background effects can obscure true biological signals in these high-throughput experiments^36^. For example, previous iterations of CiBER-seq found numerous guides whose apparent phenotype likely reflected the differing genetic requirements of distinct promoters^5^. We find that expressing RNA barcodes from closely matched promoters eliminates such background both by ensuring the reporter and normalizer respond similarly to perturbations that affect their shared genetic dependencies and reducing technical variation in sample processing. Indeed, we infer dramatically different guide activities within the same sample depending on the normalization approach. The lack of significant guide effects in the absence of differential regulation of highly similar transcription factors argues that matched promoters minimize systems-level effects and favors this design for barcode-based CRISPRi screens.

The improved quantitation and precision of this optimized CiBER-seq approach addresses a major challenge in functional genomics. In profiling regulation of the CL1 degron, we found all genes with established roles in this pathway with minimal background. Our approach also offers to expand our ability to interrogate regulators of critical biological processes by directly interrogating post-transcriptional regulation of RNA reporters. Using NMD as a model, CiBER-seq highlighted the known NMD factors and confirmed their dependence on ribosome elongation. This platform now opens opportunities to directly assess RNA-level phenotypes with CRISPR screens and paves the way to examine post-transcriptional processes that do not require translation, such as nuclear RNA decay or noncoding mRNA metabolism^37–40^.

While this work provides a general blueprint for applying CiBER-seq, additional considerations may arise in specific applications. Closely matched sample processing and reporters with similar genetic dependencies reduce noise, and these should be carefully evaluated when adapting this technique to other phenotypes. Incorporating additional reporters or perturbations can further expand the range of processes accessible to barcode-based genetic screens. Furthermore, because CiBER-seq is a sequencing-based, massively parallel reporter assay across genetic perturbations, we anticipate that it can be combined with single-cell sequencing to interrogate specific molecular processes in complex cellular environments^41^. These platforms will enable precise dissection of the intricate genetic networks controlling the processes governing cellular behaviors from the biochemical to organismal scale.

## Acknowledgements

We thank Tina Sing and Timothy J. Eisen for helpful comments on the manuscript and members of the Ingolia and Liana Lareau laboratories for suggestions throughout the study. This work was supported by NIH Ruth L. Kirschstein Postdoctoral Fellowship 5F32GM148044-02 (J.H.L) and R01 GM135233 (N.T.I). Sequencing was performed at the UCSF CAT, supported by UCSF PBBR, RRP IMIA, and NIH 1S10OD028511-01 grants.

## Author contributions

J.H.L and N.T.I designed the study. J.H.L performed all CiBER-seq experiments and analysis. N.T.I adapted the Bxb1 recombinase for use in yeast. J.H.L wrote the manuscript with input and editing from N.T.I, who supervised the project.

---

**Figure S1:**
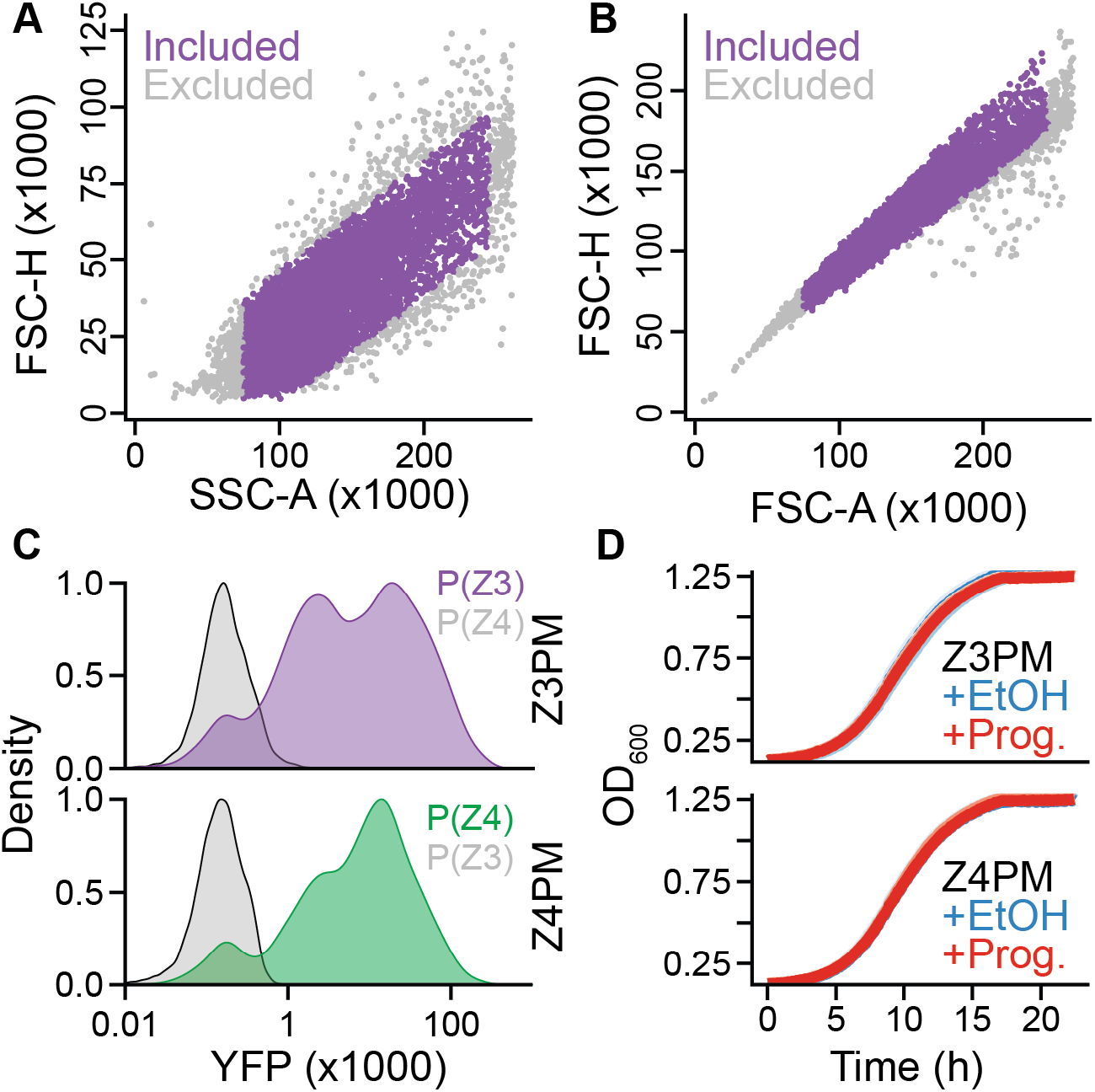
Evaluation of Z3PM and Z4PM for reporter expression. **(A-B)** Representative gating criteria for forward and side scatter in all flow cytometry analysis. **(C)** Representative raw flow cytometry histograms of Z3PM or Z4PM driving YFP expression from P(Z3) or P(Z4) at 200 nM progesterone. **(D)** Raw growth curves from yeast containing Z3PM or Z4PM and expressing YFP from the cognate promoter, with 200 nM progesterone or matched volume of ethanol in synthetic complete media lacking uracil (n=3).

**Figure S2:**
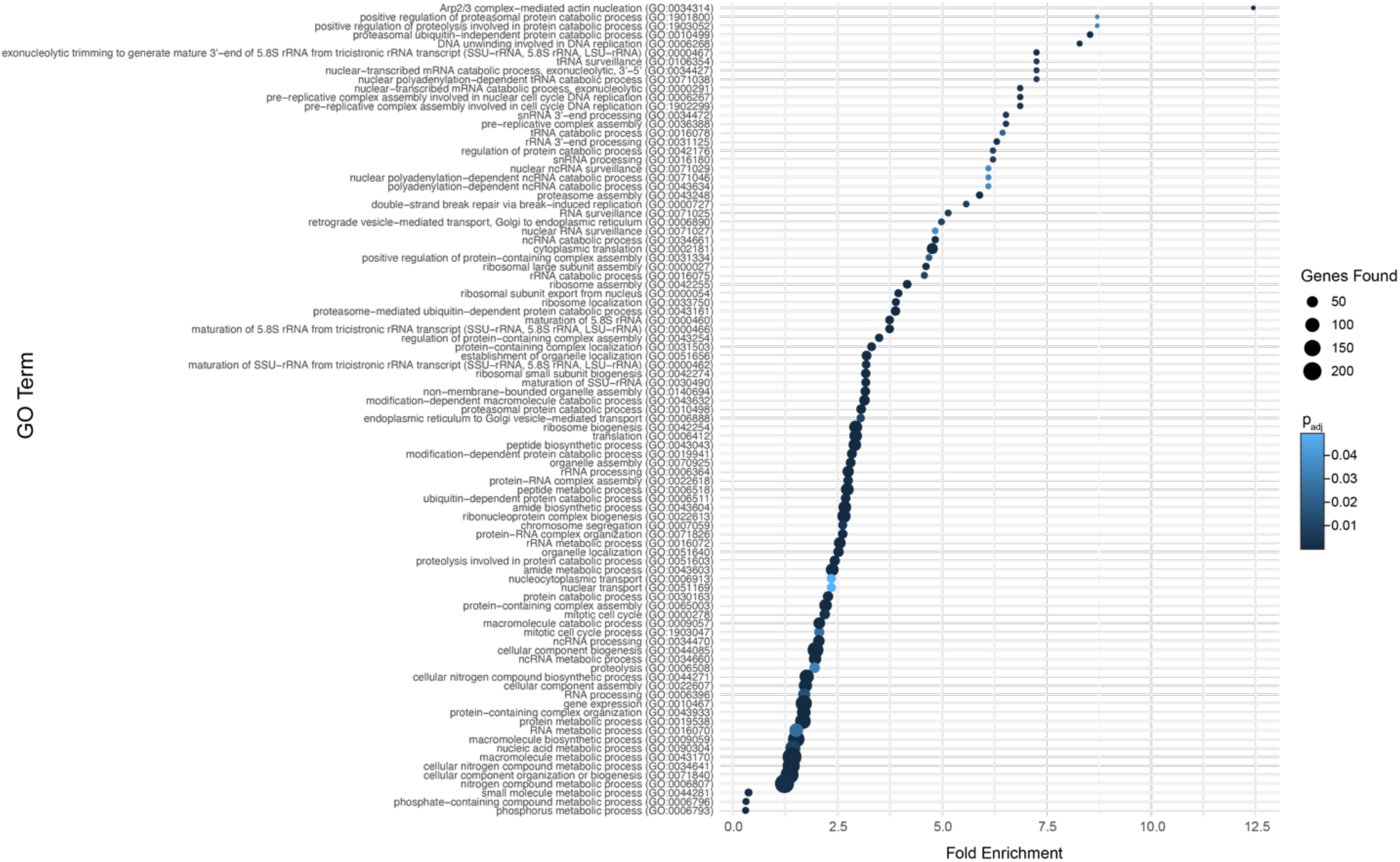
Analysis of significant processes in DNA-normalized CiBER-seq. Gene ontology terms for guides that significantly increased barcode expression in Fig. 1G, with significance (*q* < 0.05) calculated from the Fisher’s exact test with the Bonferroni correction. There were no significant gene ontology terms for guides that decreased barcode expression in the DNA to RNA comparison.

**Figure S3:**
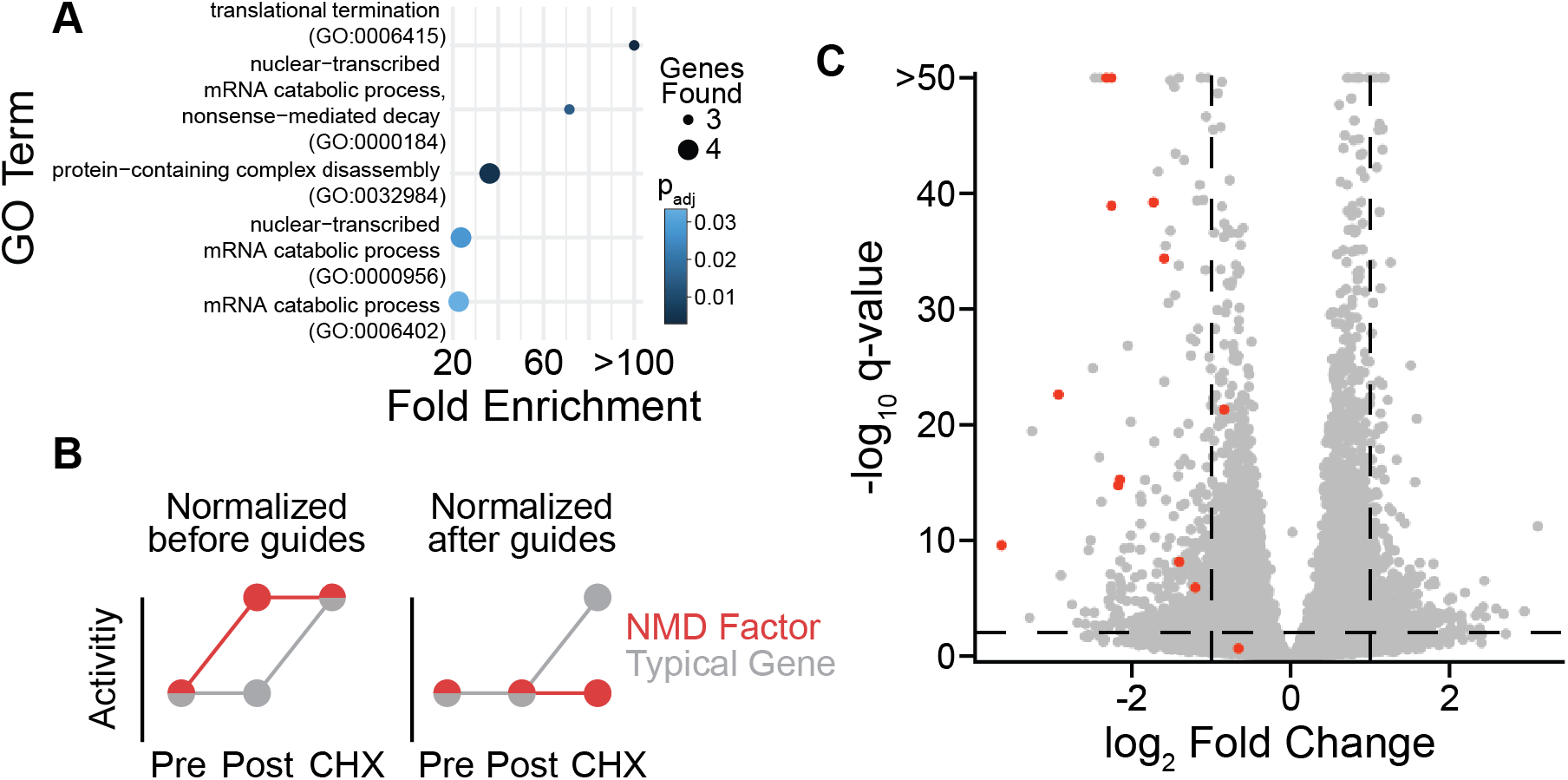
CiBER-seq profiles the requirement of activate translation for NMD activity. **(A)** Gene ontology enrichment analysis of guides that stabilize the PTC containing reporter compared to all guides from Fig. 4E. Significance (*q* < 0.05) was assessed with Fisher’s exact test with the Bonferroni correction. No significant gene ontology terms were found for guides that appear to further destabilize the reporter. **(B)** Schematic of expected guide activities based on the comparisons in linear models. NMD factors should have lower activity than a typical gene in the cycloheximide treatment, because the reporter harboring the PTC is not further stabilized by drug treatment. **(C)** Analysis of genome-wide CiBER-seq screen for NMD factors after cycloheximide treatment, normalized to post-guide induction. Guides targeting NMD factors, as determined in Fig. 4E, are labelled in red. Dashed lines represent q-values < 0.01 and > 1 log_2_-fold change.

## Materials and methods

### Yeast strain construction

All *S*.*cerevisae* strains used were derived from BY4742, grown at 30 °C, and transformed with the LiAc/PEG method as described previously^42^. Unless otherwise stated, all yeast were collected by centrifugation at 3000 g for 5 min at room temperature. Successful integration of reporter transcription factors, Cas9/TetR, or the attP landing pad was verified by PCR amplification across sequence junctions. For constructing PEST-Z3PM and PEST-Z4PM containing strains, these transcription factors were cloned into separate easyclone 2.0 vectors through Gibson assembly to make pJL200 and pJL201^43^. The CL1 degron was appended to the C-terminus of PEST-Z3PM to make pJL259. In all cases, vectors were linearized through PluTI (NEB, R0713S) digestion for 1 hr at 37 °C, transformed into log phase yeast using the LiAc/PEG approach, and selected for on the appropriate media. A complete list of strains, sequencing primers, and plasmids used in this study are available in Table S1.

### Transcription factor hormone response

Yeast expressing transcription factors and promoters were grown in SCD-Ura supplemented with the appropriate amount of progesterone (Sigma-Aldrich, P0130). Overnight cultures of yeast were backdiluted to OD_600_ = 0.1 and grown until OD_600_ = 0.5 before being collected by centrifugation, washed once with PBS, and then fixed in 4% paraformaldehyde/PBS solution at room temperature in the dark for 15-60 min. Fixed cells were pelleted, resuspend in PBS, and stored at 4 °C until ready for use. For testing transcription factor orthogonality, strains with genomically integrated Z3PM or Z4PM and plasmid containing P(Z3)-YFP or P(Z4)-YFP were grown in SCD-Ura with 200 nM progesterone as described above. Flow cytometry was performed on a BD LSRFortessa cell analyzer using the appropriate cytometer settings, gated for single cells, and analyzed using *flowCore* in R^44^. Hormone response curves and k_1/2_ were fit to the following equation in R:

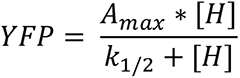

Where H is the progesterone concentration and A_max_ represents a fit variable to the maximum YFP. All measurements were derived from 2-3 biological replicates.

### Optimized CIBER reporter design and library construction

We chose to use matching P(Z3) and P(Z4) promoters that are recognized by the Z3- and Z4-DNA binding domains, respectively^10^. For protein-level phenotypes, these promoters drove expression of reporter mRNAs that encoded yeast optimized YFP and CFP^45^. The CFP transcript contained several synonymous mutations to allow specific primer binding during library amplification. For the NMD reporter, divergent *PGK1* promoters drove expression of a reporter mRNA encoding a YFP with a premature termination codon or a full-length yeast optimized mScarlet reporter^46^. In both cases, reporters were flanked by homology to the dual barcoded-guide library and BciVI sites that allowed excision of the reporters for library construction.

All CiBER screening libraries were derived from a previously reporter library containing an sgRNA paired with two unique 25 nucleotide barcodes^5^. A step-by-step description of guide cloning, barcode insertion, and barcode-guide pairing are detailed elsewhere^9^. For cloning reporters into the barcoded libraries, 2 µg of the sgRNA-barcode library was linearized with AscI (NEB, R0558S) and, separately, 2 µg of the plasmids containing the reporters and promoters (pJL206 or pJL261) were linearized with BciVI (NEB, R0596S) for 1 h at 37 °C. The digestions were purified by spin column (Qiagen, 28104) according to the manufacturer’s instructions. 480 ng of the sgRNA-barcode library and 230 ng of the reporter construct were mixed in separate 20 µL NEBuilder HiFi assembly reactions (NEB, E2621S) for 1 h at 50 °C. The assembly was then purified with the Monarch PCR & DNA cleanup kit (NEB, T1030S), eluted in 6 µL of water, and electroporated into 5 × 20 µL MegaX DH10B T1^R^ Electrocomp cells (ThermoFisher C640003) according to the manufacturer’s protocol. Cells were shaken at 37 °C for 1 h in recovery media, transferred into 200 mL LB+Kan+Cam, and grown overnight at 28 °C to OD_600_ = 3.0. Library coverage was estimated by serial dilution of transformations before overnight growth onto selective plates to ensure > 100 unique transformants for each barcode in the library. CiBER-seq libraries were then purified from overnight cultures by Midiprep (Qiagen 12143) according the manufacturer’s protocol.

### CiBER-seq screens

The reporter strains were transformed with the appropriate library by inoculating 250 mL YEPD with yeast to OD_600_ =0.1 and growing cells for 2-3 doublings. Cells were collected by centrifugation at 3000 g for 10 min at room temperature and transformed using a 30x scaled up LiAc/PEG transformation protocol with > 40 µg of the appropriate CiBER-seq library for each of two biological replicates^42^. After heat shock at 42 °C for 45 min, cells were collected by centrifugation at 3000 g for 5 min, transferred to 500 mL SCD-Ura, and grown at 22 °C for ∼2 days until OD_600_ = ∼2. After reaching the target density, 210 OD units were harvested by centrifugation and stored at -80 °C in SCD-Ura + 15% glycerol until ready for use. We estimated the number of unique transformants by serial dilution and plating on SCD-Ura, ensuring > 10x of each barcode in the library.

To perform the CiBER-seq screens, glycerol stocks were thawed and glycerol was removed by pelleting cells and washing with fresh SCD-Ura three successive times. Cells were transferred into a custom turbidostat and grown at constant turbidity (target OD_600_ = 1.0) in the appropriate media^14^. For the plasmid-based CiBER-seq screen comparing P(Z3) to P(Z4) and the CL1 CiBER screen with Z3PM-CL1 and Z4PM, cells were grown in SCD-Ura supplemented with 200 nM or 135 nM progesterone, respectively. For the NMD reporter screen, cells were grown in SCD-Ura with no hormone. Cells were grown for 16+ hours and 10 OD_600_ units were collected by centrifugation as a pre-induction sample. Anhydrotetracycline (Takara Bio, 631310) was then added to the growth chambers and the media reservoir to a final concentration of 250 ng/ml, and cells were grown for ∼9 hours before collecting 10 OD_600_ units for a post-induction sample. For the NMD CiBER-seq screen, cycloheximide (Sigma-Aldrich, C1988) was added to the media to a final concentration of 100 µg/mL and cells were grown for another hour before collecting 10 OD_600_ units. After removing the media, all samples were flash frozen in LN_2_ and stored at -80 °C until ready for library preparation. All CiBER-seq screens were performed in two biological replicates from independent library transformations.

### Barcode isolation, library preparation, and deep sequencing

RNA barcodes were isolated from cell pellets by phenol:chloroform extraction and ethanol precipitation followed by cDNA synthesis and library construction with low cycle PCR. Cell pellets were resuspended in 440 µL of 50 mM NaOAc pH 5.2, 10 mM EDTA, 1% SDS (v/v) and 400 µL phenol/chloroform and heated with agitation at 65 °C for 15 minutes before chilling on ice for 5 min. The sample was centrifugated at 20,000 g at 4 °C for 5 min, washed twice with 400 µL chloroform, and the aqueous phase was transferred to a new RNAse-free tube. RNA was precipitated by adding 50 µL of 3 M NaOAc pH 5.2, 1 µL GlycolBlue (Invitrogen, AM9515), and EtOH to 80% (v/v). The RNA was pelleted by centrifugation at 20,000 g at 4 °C for 10 min, washed once with 500 µL ice-cold 70% EtOH, and air dried for 5 minutes before resuspending in 50 µL water and quantified by nanodrop. 20 µg of RNA was treated with TURBO DNAse (Invitrogen, AM2238) in 100 µL volume for 30 min at 37 °C and then further purified by spin column (Zymo Research, R1013) according to the manufacturer’s instructions. After DNAse treatment of the cycloheximide treated samples, 10 µg of RNA was treated with 1 µL Xrn1 (NEB, M0338S) for 1 h at 37 °C to remove decapped, 5’ phosphorylated mRNAs before being purified by spin column. Following RNA isolation, 4 µg of each sample was reverse transcribed using dT priming with Protoscript II (NEB, M0368L) and subsequently treated with 0.5 µL each of Rnase H and Rnase A (Thermo Fisher, EN0531 and NEB M0297S) for 30 min at 37 °C. cDNA was purified by spin column clean up (Qiagen, 28104) and used as input for step-1 PCR with oJL923, oJL924, and oJL1058 along with oJL555 for YFP, CFP, and mScarlet amplicons, respectively. All PCR was performed in 50 µL reactions with a 95 °C initial denaturation for 3 min, followed by 7-10 cycles of 98 °C for 20 s, 60 °C for 15 s, and 72 °C for 10 s, with a final extension of 72 °C for 2 minutes using Q5 Hot Start High-Fidelity DNA Polymerase (NEB M0493L). We used 7 cycles for all amplicons, except those from the CL1-YFP sample, which were expressed at low levels and required 10 cycles. Following step-1 PCR, samples were purified by spin column and eluted in 50 µL water. Half the elution was used as input for step-2 PCR with Illumina-compatible dual index primers containing unique dual indexes (provided by UCSF CAT), using 8 cycles for the CL1-YFP sample and 7 cycles for all other transcripts. Amplicons were then purified using AMPure XP beads (Beckman Coulter, A63882) using a 1:1 bead:PCR ratio, washed once with 180 µL 80% EtOH and eluted in 15 µL water. Samples were quantified using qPCR using DyNAmo HS SYBR Green qPCR (ThermoFisher, F410L) on a Stratagene Mx300P instrument, pooled, and further analyzed by automated electrophoresis using an Agilent Tapestation 2200. Pools were sequenced on an Illumina Novaseq-X with single-end 100 bp or paired-end 150 bp reads.

For extraction of DNA barcodes, pellets were thawed and plasmids were isolated using the Zymoprep Yeast Plasmid Miniprep II kit (Zymo Research, D2004) with minor modifications: The pellet was resuspend in 1 mL of solution-1 and 30 µL zymolase was added, digesting for 3 h with agitation at 37 °C. The reaction was then split into 5 tubes and then prepared separately according to standard protocol. After pelleting precipitant, the samples were loaded successively onto a single spin column to minimize nonspecific plasmid loss and then washed and eluted according to standard protocol. YFP barcodes were first amplified for 8 cycles using oPD573 and oJL521 and then purified by spin column. Half the elution was used as input to another PCR to add overhangs for sequencing primers using oJL923 and oJL555 for 7 cycles. The sample was again purified by spin column and sequencing primers were added as described above using another 7 cycles of amplification. Samples were then purified, quantified, and analyzed as described above.

### Analysis of barcode counts and determining guide effects

Illumina sequencing reads were trimmed with cutadapt, mapped to guides using bowtie2 with the appropriate barcode sequences, and counts were extracted from alignments with samtools^47–49^. Barcodes with fewer than 5 reads in the pre-induction sample were discarded and excluded from downstream analysis. Reads for each condition were analyzed using the *mpra* package in R, summing counts for each guide across the independent barcodes^15^. Guides with a false discovery rate corrected p value < 0.01 were scored as statistically significant, and adjusted p values < 10^−50^ were set to 10^−50^ for display. All screens assessed the change in barcode counts after guide expression, using the pre-anhydrotetracycline sample as the baseline. For the cycloheximide comparison, the post-anhydrotetracycline sample was used as the baseline.

### Gene ontology analysis

Gene ontology analysis was performed using PantherDB to calculate statistical overrepresentation using the Fisher’s exact test and significance was assessed with a Bonferroni corrected p value < 0.05^50^. Full GO terms with fold and enrichment and calculated significance are displayed. We did not display comparisons that have no significant GO terms.

### Bxb1-mediated integration

Two independent clonal isolates of yeast containing the integrated Bxb1 landing pad derived from pNTI829 were each transformed with 100 fmol ARS/CEN or Bxb1 donor plasmid DNA as described above. The ARS/CEN transformation used 50 fmol (175 ng) each pNTI854 and pNTI855, while the Bxb1 donor transformation used 50 fmol (140 ng) each pNTI832 and pNTI833. A small fraction of yeast from each transformation were taken for serial dilution and plating on selective SCD-Ura media to estimate transformation efficiency. The rest of the transformation was inoculated into 50 mL pre-warmed SCD-Ura at an OD_600_ of ∼0.1 and incubated for 22 hours at 30 ºC with shaking and aeration, at which point the OD_600_ had reached 1.9 for the ARS/CEN transformation and 0.28 for the Bxb1 transformation. Yeast were then diluted into 20 mL of fresh pre-warmed SCD-Ura to achieve an estimated OD_600_ of 0.025 and growth was continued at 30 ºC with shaking and aeration for another 17 hours. A sample of yeast was fixed for flow cytometry, as above. Non-selective cultures were established by diluting yeast into pre-warmed YEPD at an OD_600_ of ∼0.1 and growing cells for an additional 6 hours with shaking and aeration, at which point they reached an OD_600_ of ∼1.5. An additional sample of yeast were fixed at this point and measured on a Benton-Dickson LSR Fortessa X20 Analyzer. yECitrine and yEmScarlet-I fluorescence was measured for 100,000 total events per sample, and gating and analysis was performed using *flowCore* in R^44^.

### RT-qPCR

Effects of individual guides were validated by performing RT-qPCR on 2-3 biological replicates of the appropriate yeast strain (Table S1). Yeast were grown overnight to saturation in SCD-Ura and then back diluted to OD_600_ = 0.1 into SCD-Ura in a 96 well deep-well block, adding 135 nM progesterone if needed. When cells reached OD_600_ = 0.5, anhydrotetracycline was added to a final concentration of 250 ng/mL and cells were grown for 16 hours. The following morning, yeast were further back diluted to OD_600_ = 0.5 and grown for 6 more hours in SCD-Ura + 250 ng/mL anhydrotetracycline, and progesterone when appropriate. For validation of individual guides, yeast were collected by centrifugation and RNA was isolated by the formamide-EDTA approach^51^. Briefly, yeast pellets were resuspended in 40 µL FAE solution (98% formamide, 10 mM EDTA), gently mixed, and heated at 70 °C for 10 minutes. Samples were then quickly vortexed, pelleted by centrifugation at 16,000 g for 2 min at room temperature, and 35 µL of the supernatant was transferred to a new tube. RNA was further purified by spin column (Zymo Research, R1018) and DNA was removed by on-column DNAse digestion according to the manufacturer’s protocol. RNA was then quantified by nanodrop and ∼500 ng of RNA was used as input for cDNA synthesis using dT priming with Protoscript II Reverse transcriptase (NEB, M0368S). cDNA was then further diluted 2-fold in water and used as input for qPCR as described above. All qPCR was performed with oJL555 with oJL923, oJL924, and oJL1058 to detect YFP, CFP, and mScarlet, respectively. Quantification of the NMD reporter was in the pre, post, and cycloheximide samples was performed on cDNA isolated from the CiBER-seq screens for the YFP and mScarlet transcripts.

### Growth curves

Yeast expressing the appropriate transcription factor and promoter combination were grown overnight in SCD-Ura and backdiluted to OD_600_ = 0.1 the following morning. Cells were grown to OD_600_ = 0.5 and then further backdiluted to OD_600_ = 0.05 in SCD-Ura with 200 nM progesterone or a matched volume of ethanol as a control. 100 µL of cells were transferred to a 96 well U-bottom plate, sealed with a Breathe-Easy gas permeable seal, and grown at 30 °C with constant agitation in a Tecan Spark plate reader while measuring the OD_600_ every 5 minutes.

## Data and code availability

Raw sequencing data and processed reads generated from this study has been deposited in the NCBI Gene Expression Omnibus (GSE268777). Code, analysis, and raw data is available on Github (https://github.com/ingolia-lab/CiBERopt_main).

